# Curcumin Alleviates Systemic Inflammation and Gut Dysbiosis Induced by Circadian Rhythm Disruption in a Rodent Model of Jet Lag

**DOI:** 10.64898/2026.01.18.698756

**Authors:** Tanya Mandyam, Michael Licamele, Martin Besmer, Gregory Peters, Sierra Simpson

## Abstract

Circadian rhythm disruption is increasingly recognized as a systemic stressor that promotes immune dysregulation and gut microbial imbalance, processes implicated in a wide range of inflammatory and neurodegenerative diseases. However, therapeutic strategies targeting the gut–immune interface under conditions of circadian misalignment remain limited. Here, we investigated whether curcumin, a plant-derived polyphenol with known anti-inflammatory properties, mitigates inflammation and gut dysbiosis induced by severe circadian disruption in a rodent model of chronic jet lag

Rats were subjected to repeated 12-hour inversions of the light–dark cycle and treated daily with curcumin (40 mg/kg/day) or vehicle delivered orally in almond butter. Circadian disruption significantly increased circulating proinflammatory cytokines and altered gut microbial composition. Curcumin treatment markedly reduced plasma levels of IFN-γ, TNF-α, IL-6, and CXCL1, decreased Peyer’s patch size, and partially restored circadian-regulated activity patterns. Shotgun metagenomic analysis revealed that curcumin shifted the gut microbiome toward a more eubiotic profile, characterized by increased species richness, reduced dominance of inflammatory taxa, decreased relative abundance of Proteobacteria, and increased Firmicutes, with a trend toward enrichment of Actinobacteria.

Collectively, these findings demonstrate that curcumin attenuates systemic and intestinal inflammation associated with circadian rhythm disruption, likely through combined suppression of proinflammatory signaling and modulation of the gut microbiome. Despite its limited systemic bioavailability, curcumin exerted robust effects at the gut–immune interface, highlighting the microbiome as a critical therapeutic target for chronobiology-associated inflammatory disorders. These results support curcumin as a potentially promising chronoprotective intervention for conditions characterized by circadian misalignment, including shift work and jet lag.

## INTRODUCTION

Inflammation is a fundamental biological response implicated in most acute and chronic debilitating diseases (Furman et al., 2019). Prolonged or unresolved acute inflammation can lead to a wide range of pathological conditions, including rheumatoid arthritis, diabetes, cancer, Alzheimer’s disease, and atherosclerosis, as well as pulmonary, autoimmune, and cardiovascular disorders, making it a major contributor to global morbidity (Ballerini et al., 2022). Inflammation is a dynamic process involving a complex network of mediators, diverse immune cell populations, and multiple metabolic pathways (Grivennikov et al., 2010). Among the key molecular players are cyclooxygenases-1 and −2 (COX-1 and COX-2) and prostaglandin-E2 (PGE2), which are central to the initiation and propagation of inflammatory responses (Smith, 1989).

Current therapeutic strategies for inflammatory diseases primarily involve steroidal and nonsteroidal anti-inflammatory drugs (NSAIDs), which act by inhibiting COX-2 and PGE2 (Ballerini et al., 2022). However, chronic use of these agents is associated with significant adverse effects, including gastrointestinal, cardiovascular, and renal complications. This underscores the urgent need to identify novel anti-inflammatory agents with improved safety profiles(Kulkarni et al., 2006). Plant-derived compounds and phytoconstituents represent a promising class of organic anti-inflammatory therapeutics (JB et al., 2003).

A growing body of research highlights the critical role of circadian rhythms in modulating inflammatory responses. These endogenous, near-24-hour oscillations regulate a wide array of physiological processes, including metabolism, immune function, and behavior. In mammals, circadian rhythms are governed by a hierarchical system of molecular clocks, with the suprachiasmatic nucleus (SCN) in the hypothalamus acting as the central pacemaker (Takahashi and Takahashi, 2016). The SCN receives photic input from intrinsically photosensitive retinal ganglion cells and synchronizes peripheral clocks throughout the body, aligning internal physiology with the external light–dark cycle (Sack, 2009). Disruption of this alignment which is commonly seen in shift work, jet lag, and exposure to artificial light can lead to circadian misalignment and has been associated with metabolic dysfunction, cognitive impairment, mood disorders, and systemic inflammation (Reilly et al., 2005; Scheer et al., 2009-3-17; Rensburg et al., 2020; Jerigova et al., 2022; Meyer et al., 2022; Cardinali et al., 2024).

One of the most compelling interfaces between circadian biology and inflammation is the gut microbiome, which is a complex and dynamic ecosystem of bacteria, archaea, viruses, and eukaryotic microbes residing primarily in the intestines. This "forgotten organ" plays a central role in digestion, immune modulation, and neuroendocrine signaling (Konturek et al., 2011; Bella et al., 2013; XC and C, 2014). The gut microbiota exhibits its own circadian rhythmicity, influenced by host feeding patterns, hormonal cues such as melatonin, and light–dark cycles. Disruption of host circadian rhythms can lead to microbial dysbiosis, characterized by a loss of beneficial taxa (e.g., *Bacteroidetes*) and an expansion of pro-inflammatory phyla such as Proteobacteria, which may compromise intestinal barrier integrity and promote systemic inflammation.

This bidirectional relationship between the circadian system and the gut microbiome is increasingly recognized as a critical component of the microbiota–gut–brain axis. Dysbiosis-induced increases in intestinal permeability can facilitate the translocation of microbial products such as lipopolysaccharide (LPS) into the bloodstream, activating Toll-like receptor 4 (TLR4) and triggering systemic inflammation. Elevated levels of proinflammatory cytokines, including interleukin-6 (IL-6) and tumor necrosis factor-alpha (TNF-α), have been shown to contribute to neuroinflammation and cognitive dysfunction (Grivennikov et al., 2010; Jerigova et al., 2022). Some gut bacteria exhibit melatonin-sensitive circadian rhythmicity, suggesting that host hormonal signals may train microbial clocks and that circadian disruption could desynchronize this finely tuned symbiosis (Paulose et al., 2016a).

Despite these insights, the precise molecular mechanisms linking circadian disruption, gut microbial alterations, and inflammatory signaling remain incompletely understood. Moreover, the downstream consequences for the central nervous system and the potential for therapeutic intervention are underexplored. Natural compounds with anti-inflammatory and microbiota-modulating properties offer a promising alternative. Curcumin, a polyphenol derived from Curcuma longa, also known as Turmeric, has demonstrated potent anti-inflammatory effects through inhibition of nuclear factor-kappa B (NF-κB) and microsomal prostaglandin E2 synthase-1 (Koeberle et al., 2009; Zhou et al., 2011) It also exhibits the ability to reshape gut microbial communities, increasing beneficial taxa and reducing pro-inflammatory species (Omosa et al., 2017; Cho et al., 2020; Wu et al., 2021; Zhu et al., 2024; Seady et al., 2025). Given its poor systemic bioavailability, the gut microbiota is increasingly viewed as a primary site of action for orally administered curcumin.

To evaluate the efficacy of curcumin in mitigating inflammation induced by circadian disruption, robust *in vivo* models are essential. Rodents, which exhibit strong circadian rhythmicity and predictable nocturnal activity patterns, provide a translational platform for studying the effects of light–dark cycle alterations on behavior, immune function, and microbiota composition (Basso and Morrell, 2017). In this study, a repeated 12-hour light–dark inversion protocol was employed to model severe circadian misalignment analogous to chronic jet lag and rotating shift work. This non-invasive paradigm enables assessment of systemic and neuroinflammatory responses in a clinically relevant context. We hypothesized that severe circadian rhythm disruption induces systemic inflammation and gut microbial dysbiosis, and that curcumin supplementation attenuates these effects. Inflammatory outcomes were assessed using plasma cytokine profiling, histological analysis of Peyer’s patches, and shotgun sequencing of the gut microbiome. This integrative approach was designed to elucidate mechanistic links between circadian misalignment, microbial ecology, and inflammatory signaling, and to evaluate curcumin as a potential chronoprotective and anti-inflammatory intervention.

## METHODS

### Animal Housing, Monitoring, Maintenance

Four-week-old male and female Long–Evans rats (Charles River Laboratories) were housed individually in cages equipped with Nalgene activity wheels (34.5 cm diameter × 9.7 cm width) and feeding hoppers. Wheel-running activity was recorded continuously using magnetic switches connected to a PC interface (Vital View, Starr Life Sciences Corp). Food and water were provided ad libitum and monitored three times weekly. All animals were 12 weeks of age at the conclusion of the study.

Experimental groups included Control (no intervention; n = 12), Vehicle (almond butter; n = 22), and Curcumin (n = 21). Subgroup analyses included ELISA (Control, n = 12; Vehicle, n = 12; Curcumin, n = 21), Peyer’s patch analysis (Control, n = 4; Vehicle, n = 12; Curcumin, n = 9), and microbiome analysis (Vehicle, n = 12; Curcumin, n = 9). All data collection and analyses were conducted in a blinded manner to minimize bias. Body weight was measured weekly over the 8-week study period using a calibrated precision scale

Rats were housed under a 12L:12D photoperiod with lights on at 9 PM and off at 9 AM PST. During weeks 5–6 and 10–11, the light cycle was inverted (lights on at 9 AM and off at 9 PM) to induce severe rhythm disturbance. The housing room was completely dark during the dark phase to prevent light interference.

### Curcumin Administration

Curcumin was administered orally via almond butter balls. Almond butter was selected for its high unsaturated fat content, which enhances curcumin bioavailability (Hocking et al., 2018). Balls were prepared by mixing almond butter and powdered rat chow in a 1:1 ratio. Curcumin was added to achieve a dose of 40 mg/kg body weight, adjusted for sex-specific average weights. Balls were refrigerated until use. During feeding, each rat was placed in a clean cage without bedding and given a diet ball. Consumption typically occurred within 40–60 seconds. Rats were returned to their original cages immediately after feeding.

### Wheel Running Data Collection

Wheel running activity was initiated on the day animals were placed in individual cages equipped with running wheels. All monitoring cables were connected and verified for data acquisition. Weekly backups of wheel running data were stored on a USB drive. At the conclusion of the study, data were extracted and converted to Excel format using Vital View software (Starr Life Sciences Corp), following the manufacturer’s instructions.

### Fecal Sample Collection and Metagenomic Analysis

Metagenomic analysis was employed to assess the genetic material of microbial communities, including bacteria and fungi, present in fecal samples. At designated time points, fecal pellets were collected directly into sterile cryovials and stored at –80°C until further processing.

Nucleic acid extraction was performed by the UC San Diego Microbiome Core using established protocols (Marotz et al., 2021). DNA was purified using the MagMAX™ Microbiome Ultra Nucleic Acid Isolation Kit (Thermo Fisher Scientific) and automated on KingFisher Flex robots. DNA quantification was conducted using a PicoGreen™ fluorescence assay. Metagenomic libraries were prepared using KAPA HyperPlus kits (Roche Diagnostics) and automated on EpMotion® liquid handlers (Eppendorf).

Sequencing was performed on the Illumina NovaSeq 6000 platform with paired-end 150 bp cycles at the Institute for Genomic Medicine, UC San Diego. Raw sequencing data were filtered for quality and demultiplexed. Bray-Curtis dissimilarity clustering analysis was conducted using QIIME2 for principal coordinates analysis (PCoA), generating biplots of overall microbiome composition. Shotgun metagenomic reads were taxonomically profiled using phylogeny-aware methods, with diversity metrics calculated at comparable taxonomic resolutions for consistency with prior microbiome studies.

### Metagenomic Data Processing and Phylogenetic Profiling

All whole-genome shotgun (WGS) sequence processing and microbial diversity analyses were conducted using QIIME2 (release 2022.8, version 2022.8.3) on an Amazon EC2 m5.24xlarge instance (96 vCPUs, 48 cores, 2 threads per core) running a Linux x86_64 platform with Python 3.8.13. Additional diversity analyses and post-processing of smaller files were performed on a personal workstation (Bolyen et al., 2019).

Shotgun metagenomic sequencing data were analyzed to characterize microbial community composition and functional potential. Sequences were processed using the Woltka pipeline (v0.1.4), which enables high-resolution phylogenetic and functional profiling of metagenomic datasets (Zhu et al., 2022).

Taxonomic classification was performed using the Web of Life release 1 (WoLr1) reference database, which includes 10,575 bacterial and archaeal genomes selected using a prototype selection algorithm based on MinHash distance matrices and genome quality control criteria. Taxonomic annotations were curated using both NCBI and GTDB systems, and a high-quality reference phylogeny was used to support phylogeny-aware analyses (Zhu et al., 2019).

Woltka was used to generate per-genome and per-gene feature tables, as well as functional profiles including KEGG pathways, KEGG orthologs (KOs), and KEGG enzymes (EZs). These outputs enabled both taxonomic and functional characterization of the microbial communities. Operational Genomic Units (OGUs) were used to provide higher resolution than traditional taxonomic units, enhancing the sensitivity of ecological and statistical analyses (Zhu et al., 2022).

### Plasma Isolation and ELISA

Trunk blood (500 ul) was collected into heparin-coated tubes and centrifuged at 3000 rpm for 5 minutes to separate plasma. Plasma was transferred to Eppendorf tubes and stored for ELISA analysis. Quantikine® high-sensitivity colorimetric sandwich ELISA kits (R&D Systems) were used to quantify levels of tumor necrosis factor alpha (TNF-α; RTA00), interferon gamma (IFN-γ; RIF00), interleukin-6 (IL-6; R6000B), cytokine-induced neutrophil chemoattractant-1 (CXCL1/CINC-1; RCN100), and interleukin-4 (IL-4; R4000).

### Histological Analysis of Peyer’s Patches

Segments of the small intestine (1.5–2 inches) were excised, rinsed with saline, and fixed in 4% paraformaldehyde for 4 days at room temperature. Samples were then transferred to a cryoprotectant solution (30% sucrose in saline). Images of Peyer’s patches were captured and stored as TIFF files for analysis using ImageJ software. Each image was coded to ensure blinding during analysis. The area of each Peyer’s patch and the corresponding intestinal segment were contoured separately, and the percent area of the patch relative to the intestinal segment was calculated to normalize for sample size variability.

### Statistical Analysis

All statistical analyses were performed using GraphPad Prism. Two-way ANOVA was used for body weight and wheel running data. One-way ANOVA was used for cytokine and Peyer’s patch comparisons. Microbiome composition was analyzed using PERMANOVA and unpaired t-tests. Statistical significance was defined as p < 0.05. Data are presented as mean ± SEM (Table 1).

**Table 1.** Summary of Behavioral, Immunological, and Microbiome Outcomes Following.

| Measure | Control | Vehicle<br>(Rhythm<br>Disturbed) | Curcumin<br>(Rhythm<br>Disturbed) | Statistical<br>Test | Significance |
| --- | --- | --- | --- | --- | --- |
| Wheel Running Output | Normal | ↓ Reduced | No change | 2-way ANOVA | p < 0.05 |
| Body Weight | Stable | Stable | Stable | 2-way ANOVA | ns |
| IFN-γ (pg/mL) | Baseline | ↑ Elevated | ↓ Reduced | 1-way ANOVA | p = 0.04 |
| TNF-α (pg/mL) | Baseline | ↑ Elevated | ↓ Reduced | 1-way ANOVA | p = 0.01 |
| IL-6 (pg/mL) | Baseline | ↑ Elevated | ↓ Reduced | 1-way ANOVA | p = 0.01 |
| CXCL1 (pg/mL) | Baseline | ↑ Elevated | ↓ Reduced | 1-way ANOVA | p < 0.001 |
| IL-4 (pg/mL) | Baseline | No change | No change | 1-way ANOVA | p = 0.02 |
| Peyer's Patch Size (% area) | Normal | ↑ Enlarged | ↓ Reduced | 1-way ANOVA | p < 0.001 |
| *Proteobacteria* (relative %) | Low | ↑ Increased | ↓ Reduced | t-test | p < 0.05 |
| *Firmicutes* (relative %) | Normal | ↓ Reduced | ↑ Increased | t-test | p < 0.05 |
| *Actinobacteria* (relative %) | Normal | No change | ↑ Trend | t-test | p = 0.1 |
| Alpha Diversity (Chao1 Index) | Normal | ↓ Reduced | ↑ Increased | t-test | p < 0.0001 |

Circadian Rhythm Disruption and Curcumin Treatment.

## RESULTS

The experimental paradigm used to induce circadian rhythm disruption is illustrated in Figure 1. This figure depicts activity monitoring using running wheels and the administration of curcumin or vehicle via almond butter–based diet balls. Figure 1 also provides a detailed experimental timeline and overview of all procedures, including behavioral testing, ELISA assays, Peyer’s patch analyses, and microbiome sequencing.

**Figure 1.**
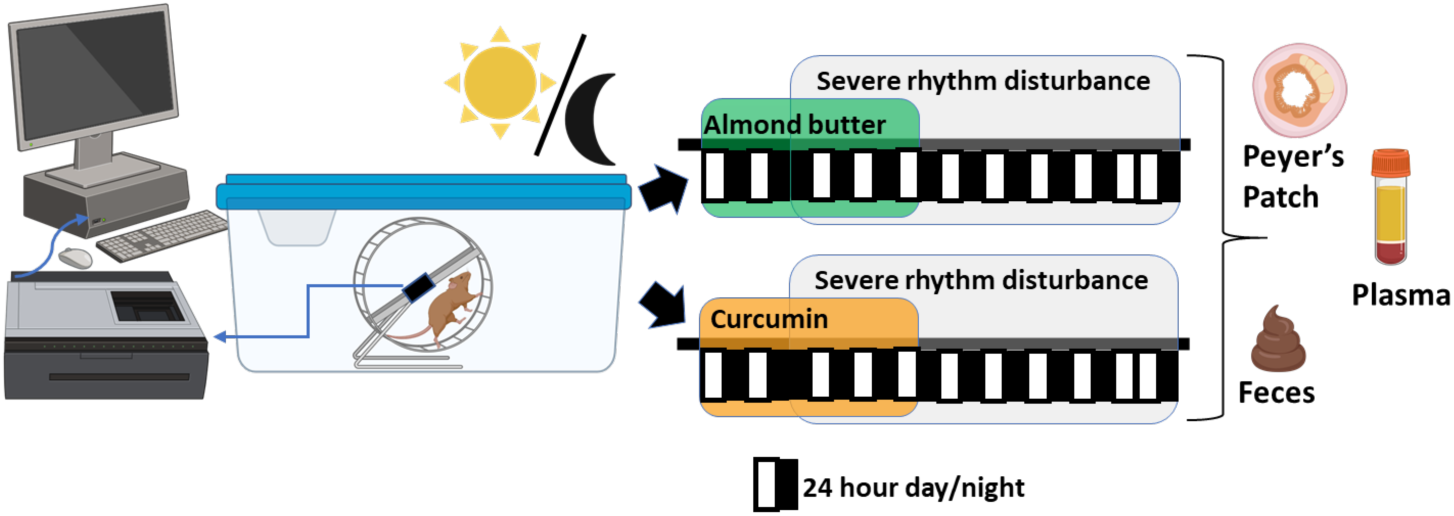
Experimental setup for inducing severe circadian rhythm disturbance in rats. Schematic in the left panel shows that animals were individually housed in cages equipped with running wheels, and wheel revolutions were monitored daily using automated software. Rats were fed either curcumin or almond butter (vehicle) diet balls. Schematic in the right panel shows the timeline and overview of experimental procedures. Following behavioral testing, blood samples were collected for ELISA-based cytokine and chemokine analysis. Small intestine segments were harvested for Peyer’s patch analysis using ImageJ. Fecal samples were collected for microbiome analysis.

### Circadian Rhythm Disruption Reduces Physical Activity

Rats subjected to a 12-hour shift in the light-dark cycle exhibited a significant reduction in wheel running activity (Figure 2). Repeated measures two-way ANOVA revealed a significant interaction between age and treatment (F_(7, 287)_ = 45, p < 0.001), with post hoc analysis showing that curcumin-treated rats maintained higher running output compared to vehicle-treated rats during the disturbance period (p < 0.05). Daily running activity before and after a 12-hour shift (day 0 to day 5) from vehicle and curcumin rats were also analyzed. Repeated measures two-way ANOVA showed a significant interaction between hourly running output x days of jet lag in both vehicle (F_(115, 2898)_ = 2.6, p < 0.001) and curcumin groups (F_(115, 2760)_ = 19.1, p < 0.001), and main effect of days in curcumin group (F _(5, 120)_ = 88.5, p < 0.001). Post hoc analysis revealed that in vehicle treated rats running pattern did not recover even beyond day 5, as running pattern was not different between days 1 to 5. However, in curcumin treated rats running pattern recovered on day 2, as running pattern was significantly different between day 1 and days 2-5 (p < 0.05). No significant differences in body weight were observed across groups.

**Figure 2.**
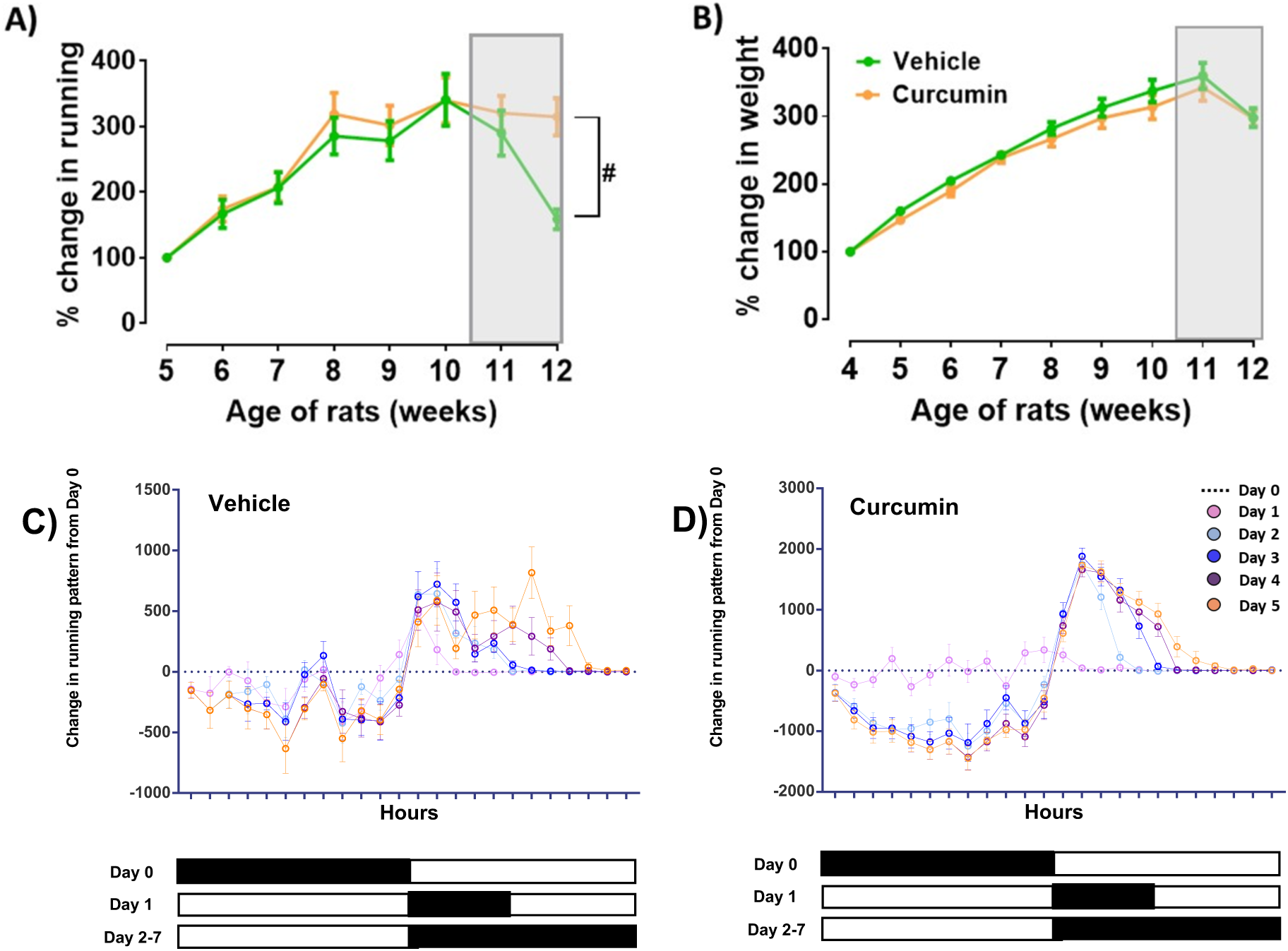
Curcumin Restores Activity Patterns Under Circadian Disruption. (A) Percent change in weekly wheel revolutions (mean ± SEM) relative to week 5 in male and female rats treated with vehicle or curcumin. The gray bar indicates the period of circadian rhythm disturbance. Two-way repeated measures ANOVA (RMANOVA, pooled across sex) revealed a significant age × treatment interaction (F _(7, 287)_ = 45, p < 0.001) and a main effect of age (p < 0.001). Post hoc Fisher’s tests showed that curcumin mitigated the reduction in running output during week 12 (p < 0.05). (B) Percent change in body weight (mean ± SEM) from weeks 5–12 relative to week 4. Two-way RM ANOVA (pooled across sex) showed a main effect of age (F _(8, 348)_ = 10.41, p < 0.001), with no significant effects of curcumin or rhythm disturbance on body weight. (C-D) Daily running activity before and after rhythm disturbance (day 0 to day 5) from vehicle (C) and curcumin (D) rats. Day 0 (normal cycle) is shown as a dotted line at 0 on the y-axis. 12-hour shift was induced on day 1, and cycle was reversed day 2-5. Two-way RMANOVA (pooled across sex) showed a significant hourly running output x days of jet lag interaction in both vehicle and curcumin groups (p < 0.05). Post hoc analysis revealed that in vehicle treated rats running pattern did not recover even beyond day 5, as running pattern was not different between days 1 to 5. However, in curcumin treated rats running pattern recovered on day 2, as running pattern was significantly different between day 1 and 2 (p < 0.05). Group Sizes: Vehicle (females): n = 8, Curcumin (females): n = 9, Vehicle (males): n = 14, Curcumin (males): n = 12.

### Curcumin Reduces Circulating Proinflammatory Cytokines

Plasma concentrations of the proinflammatory cytokines IFN γ, TNF α, IL 6, and CXCL1 were significantly elevated in vehicle treated rats following circadian rhythm disruption compared with controls (Figure 3). Curcumin treatment significantly attenuated these increases, as indicated by main effects of treatment for IFN γ (F_(2,42)_ = 3.3, p = 0.04), TNF α (F_(2,42)_ = 4.6, p = 0.01), IL-6 (F_(2,42)_ = 5.1, p = 0.01), and CXCL1 (F_(2,42)_ = 7.4, p < 0.001). In contrast, plasma levels of the anti-inflammatory cytokine IL-4 were not significantly increased by curcumin treatment, suggesting that curcumin primarily exerted its effects through suppression of proinflammatory signaling rather than enhancement of anti-inflammatory cytokine production.

**Figure 3.**
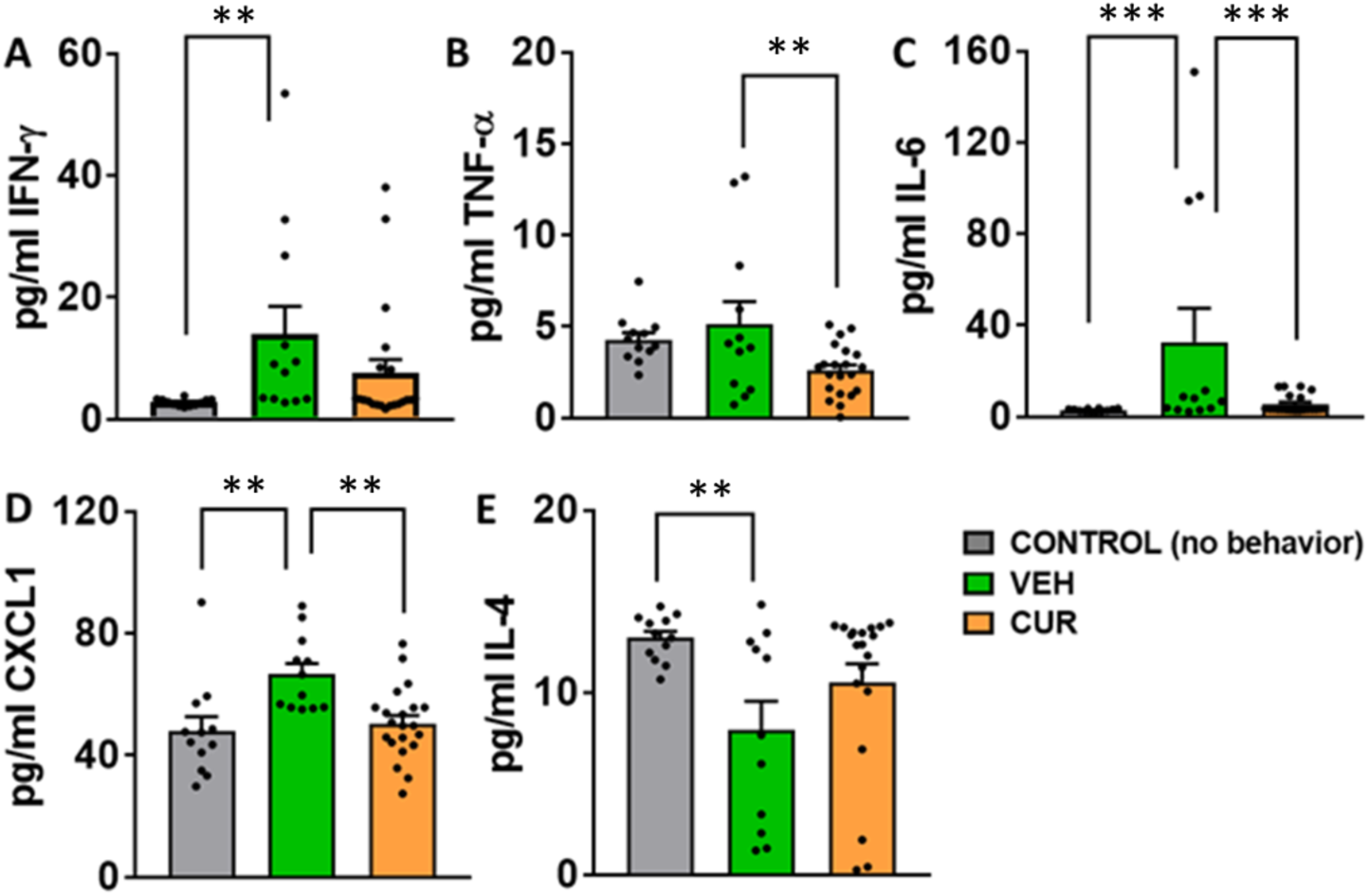
Curcumin Reduces Cytokine Elevation Caused by Circadian Rhythm Disruption. (A-E) Quantification of plasma inflammatory markers (IFN-γ, TNF-α, IL-6, CXCL1, and IL-4) in control, vehicle-treated, and curcumin-treated rats. IFN-γ, TNF-α, IL-6, and CXCL1 are proinflammatory markers; IL-4 is anti-inflammatory. One-way ANOVA revealed significant group differences for all markers: IFN-γ: F_(2, 42)_ = 3.3, p = 0.04, TNF-α: F_(2, 42)_ = 4.6, p = 0.01, IL-6: F_(2, 42)_ = 5.1, p = 0.01, CXCL1: F_(2, 42)_ = 7.4, p < 0.001, IL-4: F_(2, 42)_ = 4.2, p = 0.02. Post hoc comparisons are indicated where significant (p < 0.05). Data represent combined male and female rats.

### Curcumin Reduces Peyer’s Patch Size

One-way ANOVA revealed a significant effect of treatment on Peyer’s patch size (F _(2, 22)_ = 9.1, p < 0.001; Figure 4). Post hoc analysis showed that curcumin significantly reduced patch size compared to vehicle (p < 0.001), while no significant difference was observed between control and either treatment group.

**Figure 4.**
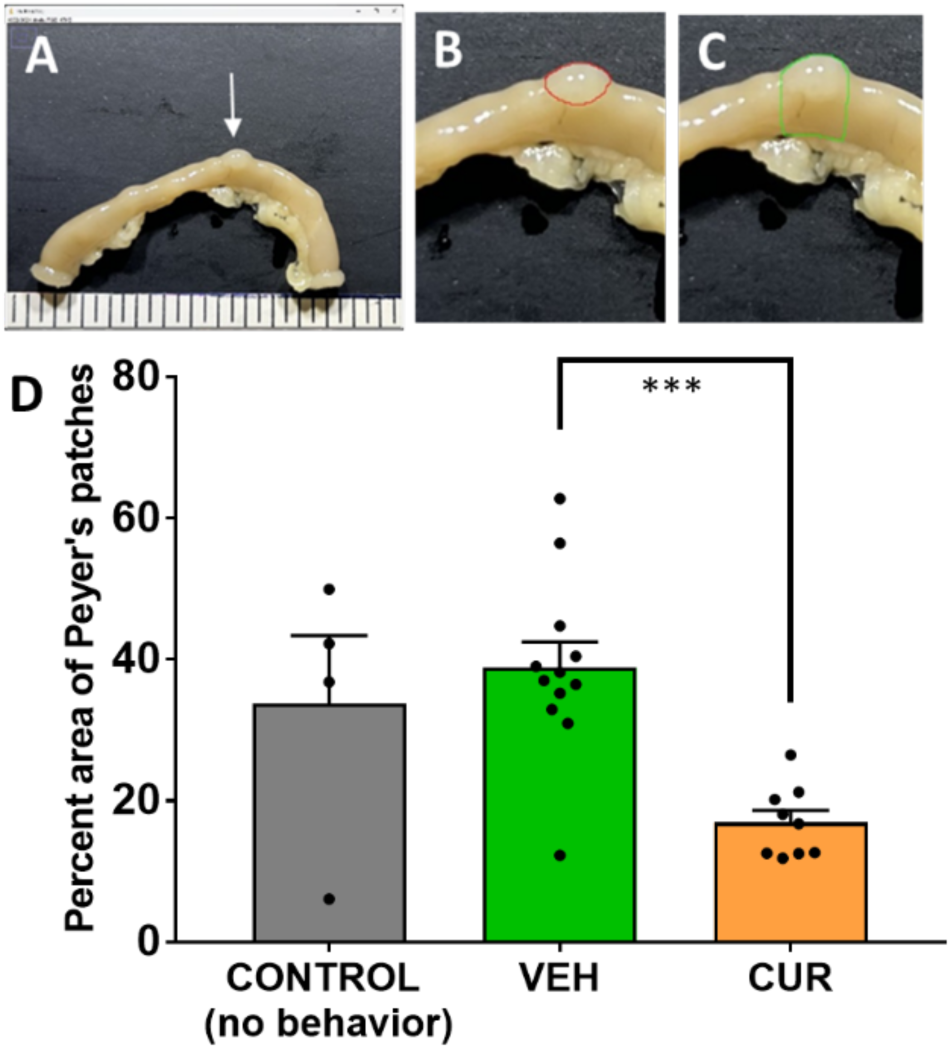
Curcumin reduces Peyer’s patch area in the small intestine. (A-C) Photomicrographs of a piece of small intestine collected and processed for Peyer’s patch analysis. Arrow in **(A)** points to a Peyer’s patch, red circle in **(B)** shows marked area of Peyer’s patch and green area in **(C)** shows the area used for total analysis. Area of Peyer’s patch was calculated as percent of area in red compared with area in green. **(D)** Quantification of Peyer’s patch size in control, vehicle-treated, and curcumin-treated rats. One-way ANOVA showed a significant group effect: F _(2, 22)_ = 9.1, p < 0.001. Post hoc analysis revealed a significant reduction in patch size in the curcumin group compared to vehicle (p < 0.001). No significant differences were observed between control and either treatment group. Group sizes: Control: n = 4; Vehicle: n = 12; Curcumin: n = 9.

### Curcumin Alters Gut Microbiome Composition

Principal coordinates analysis (PCoA) based on Bray–Curtis dissimilarity demonstrated clear separation between vehicle-treated and curcumin-treated animals, indicating a significant shift in overall microbial community composition following curcumin administration (PERMANOVA, p < 0.05; Figure 5B). Consistent with these compositional changes, alpha diversity analyses revealed a significant increase in species richness in the curcumin group, as measured by the Chao1 index (p < 0.0001; Figure 5C). In addition, microbial community structure was characterized by a reduction in dominance of highly abundant taxa in curcumin-treated animals, reflected by a lower Simpson index (p = 0.01; Figure 5D). In contrast, species evenness, assessed using the Shannon diversity index, did not differ significantly between groups (p = 0.19), suggesting that curcumin primarily influenced richness and dominance rather than overall evenness of the microbial community.)

**Figure 5.**
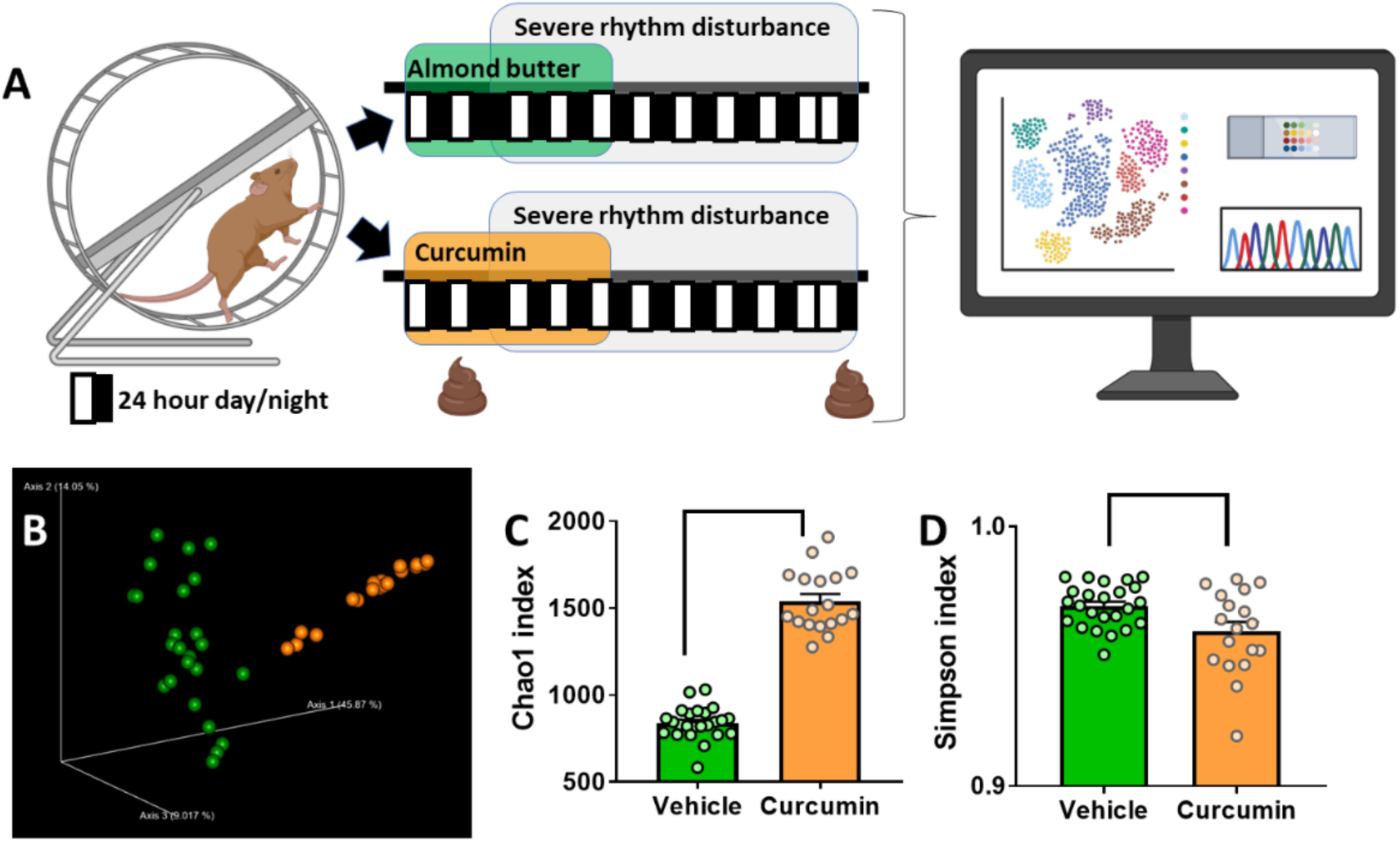
Curcumin Alters Diversity Measurements. **(A)** Experimental schematic illustrating the study design. Rats were exposed to a severe rhythm disturbance (10:10 light–dark cycles) and received either almond butter (control vehicle) or curcumin supplementation. Fecal samples were collected for 16S rRNA gene sequencing to assess microbial composition and diversity. **(B)** Principal coordinate analysis (PCoA) demonstrates distinct clustering between Vehicle (green) and Curcumin (orange) groups, indicating compositional differences in gut microbiota. **(C–D)** Alpha diversity metrics show that curcumin restored microbial richness and evenness (Shannon Index) compared to vehicle controls (mean ± SEM, p < 0.05).

### Curcumin Modulates Specific Bacterial Phyla

Taxonomic profiling revealed distinct differences in microbial composition between vehicle-and curcumin-treated animals (Figure 6). Stacked bar plots illustrate group-level shifts in relative abundance across major bacterial phyla (Figure 6A,B). A significant reduction in the relative abundance of Proteobacteria was observed in the curcumin-treated group compared with vehicle controls (p < 0.05; Figure 6C), a phylum commonly associated with intestinal inflammation and dysbiosis. In contrast, curcumin treatment was associated with a significant increase in Firmicutes abundance (p < 0.05), a phylum broadly linked to gut homeostasis and metabolic function. A trend toward increased Actinobacteria abundance was also observed in the curcumin group, although this did not reach statistical significance (p = 0.10). Collectively, these phylum-level shifts are consistent with a curcumin-associated restructuring of the gut microbiome toward a more eubiotic microbial profile.

**Figure 6.**
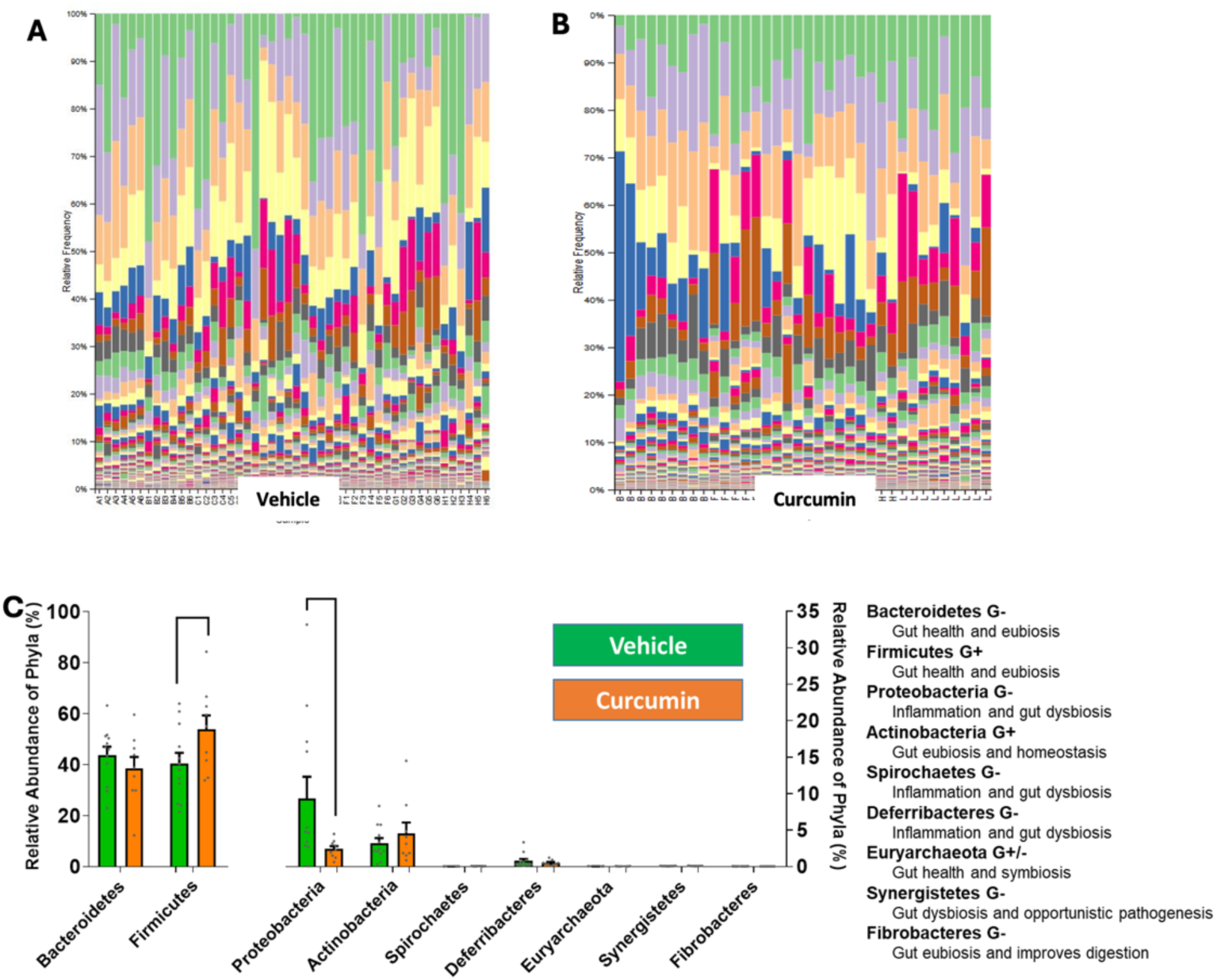
Curcumin Promotes a Shift Toward Beneficial Gut Microbiota Composition. **(A-B)** Stacked bar plots show the relative abundance of bacterial phyla in vehicle- and curcumin-treated groups. **(C)** Bar graphs showing relative abundance of the nine most abundant bacterial phyla in vehicle and curcumin groups. Phyla are classified by Gram staining characteristics (Gram-positive or Gram-negative). *Proteobacteria* was significantly higher in the vehicle group (p < 0.05). *Firmicutes* was significantly higher in the curcumin group (p < 0.05). *Actinobacteria* showed a trend toward increase in the curcumin group (p = 0.1). No other phyla showed significant differences. Group sizes: Vehicle: n = 12; Curcumin: n = 9.

## DISCUSSION

The current findings confirm that severe circadian rhythm disturbance produces systemic inflammation, consistent with previous studies in both animal models and human subjects (Jerigova et al., 2022). Inflammatory responses were evident through elevated plasma levels of proinflammatory cytokines and chemokines, including interferon gamma (IFN-γ), tumor necrosis factor alpha (TNF-α), interleukin-6 (IL-6), and cytokine-induced neutrophil chemoattractant-1 (CXCL1). These results support and extend prior work showing that rhythm disruption can activate immune pathways and contribute to inflammatory disease risk (Furman et al., 2019; Cardinali et al., 2024).

Mechanistically, inflammation involves the activation of immune cells and the release of biochemical mediators such as cytokines and chemokines (Leng et al., 2008). A key pathway includes the conversion of arachidonic acid (AA), a membrane phospholipid, into prostaglandins (PGs) by cyclooxygenase enzymes COX-1 and COX-2 (Smith, 1989; Nakanishi and Rosenberg, 2012). Among these, PGE2 plays a central role in promoting inflammation by binding to EP1–EP4 receptors on immune cells, triggering the release of inflammatory mediators (Ferrucci and Guralnik, 2003; Park et al., 2006).

The ELISA results in this study confirmed that rhythm disturbance significantly increased circulating levels of IFN-γ, TNF-α, IL-6, and CXCL1. These cytokines are implicated in a wide range of inflammatory and autoimmune diseases. IFN-γ is produced primarily by lymphocytes and is involved in both innate and adaptive immunity; elevated levels are associated with cancer and autoimmune disorders (Schroder et al., 2004; Ding et al., 2022). TNF-α, secreted by macrophages and T cells, is a potent proinflammatory cytokine linked to diabetes, atherosclerosis, and cancer (Zelová et al., 2013). IL-6, produced by monocytes and macrophages, contributes to cardiovascular dysfunction, arthritis, and Crohn’s disease(Tanaka et al., 2014). CXCL1, a chemokine that regulates immune cell migration, is associated with cancer, diabetes, and stroke (Jiang et al., 2023). Interestingly, IL-4, an anti-inflammatory cytokine produced by T cells, was not significantly increased by curcumin treatment. IL-4 is known to suppress TNF-α and IL-6 production (Velde et al., 1990; Wojdasiewicz et al., 2014), and its unchanged levels suggest that curcumin’s anti-inflammatory effects in this model were primarily due to suppression of proinflammatory pathways rather than enhancement of anti-inflammatory signaling.

In addition to systemic immune markers, this study examined Peyer’s patches which act as immune sensors in the small intestine that respond to inflammation by increasing in size (Park et al., 2023). Although rhythm disturbance did not significantly increase Peyer’s patch size compared to controls, curcumin treatment significantly reduced patch size relative to the vehicle group. This suggests a localized anti-inflammatory effect in the gut, potentially reflecting curcumin’s absorption and action in the intestinal mucosa (Meo et al., 2019).

The gastrointestinal tract is under circadian control, with feeding and fasting cycles influencing nutrient absorption, metabolism, and immune regulation (Reppert et al., 2002). Disruption of circadian rhythm can impair these processes and contribute to systemic inflammation. The gut microbiome plays a central role in maintaining homeostasis, supporting digestion, modulating immune responses, and protecting against pathogens (Robinson et al., 2010; Lozupone et al., 2012). Dysbiosis is associated with a wide range of metabolic and inflammatory diseases.

Four dominant bacterial phyla were identified: *Firmicutes*, *Bacteroidetes*, *Proteobacteria*, and *Actinobacteria* (Zo*etendal et al., 2008)*. *Firmicutes* and *Bacteroidetes* are generally associated with gut health, as they ferment dietary fibers into short-chain fatty acids (SCFAs), which serve as energy sources for intestinal epithelial cells and modulate inflammation (Wexler, 2007; Qin et al., 2010; Rinninella et al., 2019; Zafar and Jr, 2021; Shin et al., 2024). *Actinobacteria*, though less abundant, contribute to eubiosis through SCFA production and antimicrobial activity (Binda et al., 2018). In contrast, *Proteobacteria* are often associated with dysbiosis and inflammation, and their overrepresentation is considered a microbial signature of disease (Shin et al., 2015; Rizzatti et al., 2017).

In this study, severe rhythm disturbance increased the relative abundance of *Proteobacteria*, consistent with previous findings in rats exposed to light-dark cycle shifts(Voigt et al., 2014). Importantly, curcumin supplementation reversed this effect, significantly reducing *Proteobacteria* and increasing *Firmicutes*. A trend toward increased *Actinobacteria* was also observed. These shifts suggest that curcumin promotes a eubiotic microbiome, which may contribute to its anti-inflammatory effects.

Alpha diversity analyses further supported this conclusion. Curcumin-treated animals exhibited increased species richness (Chao1 index) and reduced dominance of specific taxa (Simpson index), indicating a more balanced microbial ecosystem. These findings align with recent studies showing that curcumin enhances microbial diversity and SCFA production in piglets and mice (Li et al., 2021; Recharla et al., 2021). The bioinformatics workflow and supporting diversity metrics are detailed in Supplemental Figure 1, which includes the QIIME2 pipeline and PCoA biplots showing distinct clustering between treatment groups. Supplemental Figure 2 further illustrates the experimental timeline for fecal sample collection and confirms increased species richness and reduced dominance in curcumin-treated animals.

Beyond compositional changes, circadian entrainment of the gut microbiome is an emerging area of interest. Gut bacteria exhibit diurnal oscillations in abundance and function, influenced by host feeding rhythms, melatonin signaling, and light–dark cycles (Paulose et al., 2016b). Disruption of these microbial clocks may exacerbate dysbiosis and inflammatory signaling. Although not directly assessed in this study, curcumin may influence microbial rhythmicity through its interaction with melatonin-sensitive taxa or by stabilizing feeding-driven microbial oscillations. This study was conducted in a rodent model, which may not fully capture human circadian and microbiome dynamics. Additionally, time-of-day effects of curcumin administration were not systematically evaluated, and microbial rhythmicity was inferred rather than directly measured. Future studies incorporating time-series sampling and microbial transcriptomics could clarify whether curcumin acts as a chronobiotic agent within the gut ecosystem.

Additionally, the timing of curcumin administration may critically influence its efficacy through mechanisms of chronopharmacology. Intestinal permeability, immune cell trafficking, and microbial metabolic activity all exhibit robust circadian regulation, suggesting that the anti-inflammatory and microbiota modulating effects of curcumin may vary depending on the time of administration. Aligning curcumin delivery with host circadian phases or periods of heightened microbial activity may therefore enhance its therapeutic impact and warrants further investigation.

Curcumin’s poor bioavailability has historically limited its therapeutic use (Stohs et al., 2020). However, co-administration with fat-rich foods, such as almond butter, has been shown to enhance absorption(Hocking et al., 2018). In this study, curcumin was delivered in almond butter balls, which likely improved its intestinal uptake and allowed for localized action in the gut. The observed reduction in Peyer’s patch size and modulation of gut microbiota support this mechanism.

The proposed model of curcumin’s action involves inhibition of PGE2 synthase-1, reducing PGE2-mediated cytokine release(Koeberle et al., 2009), and enhancement of SCFA-producing bacteria, which further suppress COX-2 activity and inflammation (Burapan et al., 2017; Liu et al., 2020). This dual mechanism is visually summarized in Supplemental Figure 3, which illustrates curcumin’s inhibition of PGE2 synthase-1 and its role in promoting SCFA-producing bacteria. These dual mechanisms involving direct inhibition of inflammatory signaling and microbiome mediated modulation may explain the broad anti-inflammatory effects of curcumin observed in this study.

This study provides compelling evidence that curcumin supplementation can alleviate systemic inflammation and gut dysbiosis induced by severe circadian rhythm disruption. Using a multi-modal approach that includes behavioral monitoring, cytokine profiling, histological analysis, and shotgun metagenomic sequencing, we demonstrated that curcumin significantly reduced circulating levels of proinflammatory cytokines (TNF-α, IL-6, CXCL1, IFN-γ), decreased Peyer’s patch size, and promoted a eubiotic gut microbiome by reducing *Proteobacteria* and increasing *Firmicutes* and *Actinobacteria*. These effects are likely mediated through dual mechanisms: inhibition of PGE2 synthase-1 and enhancement of SCFA-producing microbial populations(Koeberle et al., 2009; Burapan et al., 2017; Liu et al., 2020).

These findings reinforce the concept that circadian rhythm disruption is not merely a behavioral or neurological disturbance, but a systemic insult with immunological and microbial consequences. Circadian misalignment, which is common in shift work, jet lag, and modern light exposure patterns can desynchronize host-microbe interactions, leading to microbial dysbiosis, increased intestinal permeability, and systemic inflammation. The gut microbiome, far from being a passive passenger, emerges as a dynamic and sensitive transducer of circadian signals, capable of influencing host physiology through microbial metabolites, immune modulation, and neuroendocrine signaling.

Although conducted in a rodent model, the results have significant translational implications. Human microbiome studies show similar microbial profiles, with Firmicutes and Bacteroidetes dominating the gut and Proteobacteria and Actinobacteria present in smaller but functionally important proportions (Turnbaugh et al., 2007). Clinical studies have linked *Proteobacteria* overgrowth to inflammatory bowel disease, metabolic disorders, respiratory diseases, and certain cancers, all conditions that are characterized by systemic inflammation (Rizzatti et al., 2017). Moreover, curcumin’s protective effects on circadian-regulated behavior and gut health may be relevant for individuals experiencing sleep disorders such as insomnia, narcolepsy, sleep apnea, and Restless Legs Syndrome (Hauri, 2011). Importantly, curcumin’s efficacy in this model, despite its poor systemic bioavailability, suggests that its primary site of action is within the intestinal lumen. By stabilizing the gut microbiota and reinforcing mucosal immune homeostasis, curcumin may interrupt the pathological cascade linking circadian disruption to systemic and neuroinflammation. These findings position the gut microbiome as a tractable therapeutic interface for chronoprotective interventions.

Future research should explore curcumin’s clinical utility in human populations. Multiple Phase 0, I, and II trials have already tested high-dose curcumin (3,000–8,000 mg/day) for various inflammatory conditions with minimal side effects (Cheng et al., 2001; Epelbaum et al., 2010; Toden et al., 2015; Mirzaei et al., 2018; Pivari et al., 2019; Stohs et al., 2020; Kristopher et al., 2021). Based on the current findings, future trials should evaluate curcumin co-administered with fat-rich foods (e.g., nut spreads) to enhance bioavailability (Hocking et al., 2018). Additionally, longitudinal studies incorporating wearable circadian tracking, microbiome profiling, and inflammatory biomarkers will be essential to validate these findings and optimize dosing strategies. Mechanistic studies using fecal microbiota transplantation, gnotobiotic models, and neural circuit mapping will be critical to delineate the causal pathways linking microbial dysbiosis to neuroinflammation. Investigating the role of host-to-microbe signaling (e.g., melatonin rhythms) and gut-to-brain communication (e.g., vagal vs. humoral pathways) will further clarify the bidirectional nature of the circadian–gut–brain axis.

In summary, this study highlights curcumin as a promising, safe, and accessible intervention for mitigating inflammation and microbiome imbalance associated with circadian rhythm disruption and related disease states. As our society continues to operate on a 24/7 schedule, such integrative strategies may prove essential for preventing a wide spectrum of chronic inflammatory and neurodegenerative diseases.

## ACKNOWLEDGEMENTS

The authors thank the staff of the UC San Diego Microbiome Core and the Institute for Genomic Medicine for their technical support in sequencing and data processing. We are also grateful to Rajitha Narreddy at the VA San Diego vivarium for their assistance with animal husbandry throughout the study.

## AUTHOR CONTRIBUTIONS

Tanya Mandyam contributed to conceptualization, investigation, methodology, formal analysis, data curation, visualization, and drafting of the manuscript. Michael Licamele and Martin Besmer contributed to data analysis and manuscript preparation. Gregory Peters contributed to software development, data curation, formal analysis, visualization, and manuscript preparation. Sierra Simpson contributed to conceptualization, project administration, funding acquisition, supervision, data curation, formal analysis, visualization and manuscript review and editing, and served as corresponding author.

## ETHICAL CONSIDERATIONS

All animal procedures were approved by the Institutional Animal Care and Use Committee (IACUC) at the VA San Diego and conducted in accordance with institutional guidelines and relevant national regulations. Every effort was made to minimize animal suffering and to reduce the number of animals used.

## CONSENT TO PARTICIPATE

Not applicable. This study did not involve human participants.

## CONSENT FOR PUBLICATION

Not applicable. This manuscript does not contain any individual person’s data in any form, including identifiable details, images, or videos.

## DATA AVAILABILITY

Requests for access to anonymized data or sequencing results may be considered upon reasonable request to the corresponding author.

